# Gui Shao Di Huang Wan promotes angiogenesis and regulates oestrogen receptor α and progesterone receptor β in *in vitro* bioassays

**DOI:** 10.1101/2021.06.14.448215

**Authors:** Rebecca O’Cleirigh, Roslyn Gibbs

## Abstract

**Background and aim:** The formula Gui Shao Di Huang Wan (GSDW) is used frequently to treat female infertility. This study aims to investigate some of the possible mechanisms of action of GSDW using *in vitro* bioassays of angiogenesis in Human Uterine Microvascular Endothelial Cells (HUVEC) and Human Uterine Microvascular Endothelial Cells (HUtMEC) and ovarian steroid receptor expression in a human endometrial cell line (Ishikawa).

**Experimental procedure:** Aqueous extracts of GSDW and its component herbs were tested for pro-angiogenic activity using both HUVEC and HUtMEC 2D differentiation assays performed on Matrigel and effects on HUVEC proliferation using the MTT assay. Effects on the expression of Estrogen Receptor α (ERα) and Progesterone Receptor β (PRβ) in Ishikawa cells were determined using immunoblotting.

**Results and Conclusion:** All analysed parameters of differentiation were increased by GSDW in both the HUVEC and HUtMEC mesh. Furthermore, measures of total length, segment number, junction number were affected by some but not all component herbs.

The MTT assay showed an increase in proliferation of HUVECs at concentrations of GSDW between 0.68 and 5.47 μg/mL at 48 and 72 hours.

In Ishikawa cells downregulation of ERα and upregulation of PRβ was seen after 48 hours incubation with 4 of the 8 herbs in the formula.

The findings in this study demonstrate that GSDW has the potential to affect key parameters (vascular, sex hormone receptor expression) *in vitro*. This offers a mechanism by which these herbs may enhance fertility through improved endometrial receptivity and pregnancy rates.

## 1. Introduction

Infertility is a growing worldwide problem exacerbated by increased maternal age, high BMI, poor diet, and stressful or sedentary lifestyles. All these factors influence the endocrine system, immunity, oocyte quality, and the receptivity of the uterine environment ^1^. IVF is generally successful in oocyte retrieval, fertilisation and the production of embryos for transfer but the average live birth rate following IVF for women under 35 is around 30%. This varies between clinics, underlying pathologies, and interventions. Even using the newest technologies of non-intrusive monitoring, genetic selection, and enhanced growing media with a visually or genetically suitable embryo the success rates remain at 30-60% ^2^.

When oocyte quality is acceptable then IVF failures may be attributable to the implantation environment. If the endometrium is persistently thin, there may be recurrent implantation failure or repeated early loss. In some cases this will be due to immune complications, but a key aspect of the receptivity of the endometrium to implantation is uterine perfusion and blood vessel development ^3^. The thickness of the endometrium as measured by ultrasound has been correlated with increased embryo implantation ^4^ and the endometrium must have adequate vasculature for invasion of the blastocyst ^5^.

The menstrual cycle begins with regeneration of the functionalis of the endometrium through the proliferation of epithelial cells, accompanied by changes in the supporting endothelial cells necessary for vascularisation ^6^. During this proliferative phase oestrogen receptors numbers increase ^6^. Differentiation of endothelial cells occurs in the endometrium during initial repair in the early proliferative phase, in the mid proliferative phase, and during the formation of the spiral arteries during the early secretory phase ^7^. The increase in vascular permeability that makes the endometrium receptive to implantation is controlled by PRs. This increased permeability is vital for the proteins contained in the blood to pass into the interstitial space to support the developing blastocyst ^6^. At a simplistic level, rising oestrogen levels in the secretory phase predominantly enhances endometrial cell proliferation, while progesterone in the secretory phase leads to differentiation ^7^. Subendometrial vascularity is reduced in women with infertility ^9^ as is uterine perfusion ^10^. Increased uterine perfusion and vascular permeability enhances nutrient transport to the embryo ^11^ and facilitates the embryo maternal signalling which is necessary for successful implantation ^12^.

Clinical studies have shown that plant extracts are able to improve pregnancy outcomes in natural and assisted conception ^8–11^. According to Traditional Chinese Medicine (TCM) theory, the main causes of infertility are Kidney and Liver yin deficiency, Blood deficiency and stasis ^12^. Gui Shao Di Huang Wan is a Kidney and Liver yin, and Blood nourishing formula. It is often used as a base formula in practice and it is a classic combination frequently cited in literature on treating infertility ^13 14^. A meta-analysis of Chinese herbal medicine and fertility^9^ conducted in Australia in 2015 included 1851 women with a range of fertility disorders. The analysis found that pregnancy was three and a half times more likely using Chinese herbs and pharmaceuticals over four months than pharmaceuticals alone. Another second study measured markers of endometrial receptivity in women receiving herbal treatment for whom IVF had previously failed. Patients were treated using a four-week, four formula phased approach; in week two an unmodified Gui Shao Di Huang Wan was used and in weeks three and four, six of the eight ingredients were used in a modified formula with additional herbs. All six measured markers of endometrial receptivity were significantly improved and there was a 34% pregnancy outcome ^15^. Despite the documented evidence, the biological mechanisms that mediate the actions of these herbs are largely unknown and this lack of mechanistic knowledge can influence physicians to advise women against the use of non-pharmaceutical medication whist undergoing assisted reproduction ^16^. This study aims to investigate those mechanisms related to angiogenesis, the proliferation and differentiation of endothelial cells, and the regulation of oestrogen and progesterone receptors in an epithelial cell model to some of the commonly used Chinese herbs.

## 2. Materials and methods

### 2.1 Preparation of the herb extract

Dried herbs as listed in Table 1 were obtained from Avicenna, Hove; UK. Samples were submitted for macroscopic identification and species validation to the medical botanist at Kew Gardens Jodrell laboratory ^18^. They were verified as being consistent with the authenticated samples held for reference. An aqueous extract was prepared as described in the Pharmacopeia of the People’s Republic of China ^17^. The herbs, individually and as a complete formula, were soaked for 12h in 100mL distilled water. They were then heated to 100°C using a hot plate (Fisher Scientific / 11-600-49SH) and simmered for 20 min. The extracts were filtered and centrifuged at 3500 rpm (2383g Heraeus Labofuge 400) for 10 min to remove plant residue. The supernatant was passed through a 22μm syringe driven filter (Millex) in a Biological Class II laminar airflow cabinet into a sterile universal container (Thermo scientific) and stored at 4 °C until required. To measure the final concentration (mg/mL) of each extract and facilitate comparison between assays, 1 mL aliquots of each extract were evaporated to dryness and the dry weight of the residual (dissolved) compounds per mL determined.

**Table 1.** Herb names and grams used in preparation of the extract

| **Pharmaceutical name <sup>17</sup>** | **Scientific name** | **Chinese** | **Pinyin** | **Grams** |
| --- | --- | --- | --- | --- |
| *Radix Rehmanniae Preparata* | *Rehmannia glutinosa* (Gaertner) De Candolle | 熟地黄 | Shu Di Huang | 8 |
| *Fructus Corni* | *Cornus officinalis* Siebold & Zuccarini | 山茱萸 | Shan Zhu Yu | 4 |
| *Rhizoma Dioscoreae* | *Dioscorea oppositifolia* Carolus Linnaeus | 山药 | Shan Yao | 4 |
| *Sclerotium Poria Cocos* | *Wolfiporia extensa* (Peck) Ginns | 茯苓 | Fu Ling | 3 |
| *Rhizoma Alismatis* | *Alisma plantago-aquatica* subsp. *orientale* (Sam.) Sam | 泽泻 | Ze Xie | 3 |
| *Cortex Moutan* | *Paeonia ostii* T. Hong & J. X. Zhang | 牡丹皮 | Mu Dan Pi | 3 |
| *Radix Angelicae Sinensis* | *Angelica sinensis* (Oliv.) Diels | 当归 | Dang Gui | 4 |
| *Radix Paeoniae Alba* | *Paeonia lactiflora* Pallas | 白芍 | Bai Shao | 4 |

### 2.2 Cell culture

In this study Human Umbilical Vein Endothelial Cells (HUVEC) were chosen as a relatively robust, stable primary cell line which is well suited and extensively used for the chosen *in vitro* techniques of investigating angiogenesis. The HUVECs are macrovascular endothelial cells and so they are not directly comparable to the uterine endothelial cells which are microvascular. Endothelial cells are not heterogenous ^19^, in different parts of the vasculature they are responsible for angiogenesis, permeability, and vessel tone. It is their heterogeneity that permits this range of roles ^20^.

Human Uterine Microvascular Endothelial Cells (HUtMEC) are also a primary cell line and were also used for the angiogenesis assays. Obtained from the myometrium these cells are known to modulate adhesion molecule expression, show increased proliferation, cell survival and angiogenic activity in response to the ovarian steroids^21^.

The Ishikawa is a transformed epithelial cell line and was used to examine the effects of the herb extracts on expression of receptor proteins using immunoblotting. Although Ishikawa cells are clones of a human endometrial adenocarcinoma, their low grade (highly differentiated) phenotype and expression of both oestrogen and progesterone receptors ^22^ has led to their extensive use as a model of steroid receptor expression in reproductive biology research. Ishikawa cells were therefore chosen to examine the effects of the herb extracts on steroid receptor expression as a model of the premenopausal endometrium ^23^.

All cells were cultured in lysine coated cell culture flasks (Thermo scientific Nucleon delta EasYFlask 25cm2) and incubated at 37°C and 5% CO__2__ (Nuaire, DH Autoflow). Media was changed every 48 hours and cells passaged when they reached confluence as recommended by the supplier. Table 2 summarises the culture conditions employed for each cell line.

**Table 2.** Cells used and their cultivation specifics

| Cell type | Supplier | Medium | Supplements | Seeding density | Considered confluent | Ratio for splitting cells |
| --- | --- | --- | --- | --- | --- | --- |
| Human Umbilical Vein Endothelial Cells (HUVEC) | Caltag ZHC-2301 | Cellworks Human Large Vessel Endothelial Cell Basal Medium KC1015 | Cellworks Human Large Vessel Endothelial Cell Growth Supplement KC1016<br>Cellworks Antibiotic Supplement (Gentamycin/ Amphotericin B) KC1019 | 2,500 cells/cm<sup>2</sup> | 60-80% | 1:2 |
| Human Uterine Microvascular Endothelial Cells (HUtMEC) | 2bScientific | Promo cell Endothelial Cell Basal Medium MV2 (prf) | + Foetal Calf Serum, Epidermal Growth Factor, Basic Fibroblast Growth Factor, Insulin-like Growth Factor, Vascular Endothelial Growth Factor 165, Ascorbic Acid, Hydrocortisone | 10,000-20,000 cells/cm<sup>2</sup> | >70 % | 1:2 |
| Ishikawa | Sigma Aldrich 99040201 (from the European collection of authenticated cell cultures) | Ishikawa Culture Medium MEM | + 2mM Glutamine + 1% v/v Non-Essential Amino Acids (NEAA) + 5% v/v Foetal Bovine Serum (FBS) | 20,000-30,000 cells/cm<sup>2</sup> | 70-80 % | 1:5 |

### 2.3 Endothelial cell proliferation

The effect of GSDW on the proliferation of HUVECs was assessed using the MTT (3- (4,5-dimethylthiazol-2-yl)-2,5-diphenyltetrazolium bromide) assay ^24^. The MTT assay is commonly accepted to be a measure of proliferation, but specifically measures cell viability ^25^. It is possible that there is no increase in the number of cells but an upregulation of their mitochondrial activity. It is generally considered in most cell populations that total mitochondrial activity is related to the number of viable cells ^26^ and so is indicative of a proliferative effect. A second assay, a MultiTox assay (Promega G9270) was also performed which measures live cell protease activity. HUVECs, cultured in Cellworks Human Large Vessel Endothelial Cell Basal Medium KC1015, were seeded at 2 = 10^5^/mL in 96 well plates (Thermo scientific 167006 Nunclon delta surface) and incubated for 24 h at 37°C and 5% CO__2__ (Nuaire, DH Autoflow). Serial dilutions of GSDW from the neat decoction, concentration 86.70μg/ mL, to a 1:512 dilution of 0.17μg/mL were added to the cells and tested against a water only (no herb extract) control. Following incubation for a further 24 - 72h, cell proliferation was assessed using the MTT assay. Absorbance values measured at 560 nm were used to determine % viability of HUVEC cells following treatment with GSDW relative to the control (representing 100% viability). Due to the long doubling time of HUtMECs, it was not possible to obtain sufficient cells for viability assays as a comparator.

### 2.4 Angiogenesis differentiation assay

25 μL of HUVECs suspended in Cellworks Human Large Vessel Endothelial Cell Basal Medium containing 0.1% FBS (v/v)or 25 μL of HUtMECs (2bScientific) suspended in Promo cell Endothelial Cell Basal Medium MV2 (prf) containing 0.1% FBS (v/v) at a concentration of 3 = 10^5^ /mL, were seeded onto aliquots of 10μL growth factor deficient, phenol red free matrigel (Corning) in an 18 well μ-slide (Ibidi). Each herb extract was diluted 1:250 with cell culture media for the HUVEC, 1:500 for the HUtMEC and 25ml then added to the u-slide wells to give a final herb extract dilution of 1:500 or 1:1000. The slides were incubated at 37°C and 5% CO__2__ (Nuaire, DH Autoflow). Cell differentiation was documented using a Nikon eclipse 80i microscope coupled to a digital camera (Nikon digital sight DS-Fi2) at 4X magnification and slides were photographed at 18h for analysis of the number of junctions, number of segments, total length and mesh area using ImageJ software.

### 2.5 Protein isolation and Immunoblotting

3mL Ishikawa cells were seeded at 1×10^5^ /mL in 6 well plates (Corning costar 3516 culture cluster flat bottom plate) in media (Gibco Minimum Essential Medium) containing 5% (v/v) FCS and incubated overnight in a humidified atmosphere at 37°C and 5% CO__2__ (Nuaire, DH Autoflow). After 24 hours 1.5mL of the media was removed and media with herb extract or media only control was added. The herb extracts were diluted 1:50 with media for a final concentration in the wells of 1:100. Cells were incubated for 48 h. Pierce RIPA buffer (Thermo Scientific) was used to prepare total cell protein extracts as described in the product information sheet. Extracts were stored at −80°C. SDS-PAGE was performed using a 12.5% resolving gel prepared according to a modified Laemmli protocol ^27^ in a Bio-Rad mini protean tetra hand cast system. Separated proteins were transferred to a PVDF membrane, which was blocked with 5% (w/v) skimmed milk powder in PBS and probed for ERα at a concentration of 1:1000 (Abcam ab108398) and PRβ at a concentration of 1:1000 (Abcam ab46535) expression. Antibodies were tagged with a goat anti rabbit (GxR) secondary antibody (Abcam) at a concentration of 1:2000 and visualised using Luminata HRP substrate on a ChemiDoc MP imaging system. ImageLab was used to measure signal intensity and calculate the abundance.

## 3. Results

### 3.1 Endothelial cell – Proliferation

HUVECs at 24, 48, and 72 hours were exposed to serial dilutions of the neat GSDW starting at a concentration of 86.70 μg/mL. There was a cytotoxic effect at higher concentrations of the extract (10.84 - 86.70 μg/mL), however, at lower concentrations a dose dependent proliferative effect was observed. At concentrations below 5.42 μg/mL, the % cell viability appeared to increase above 100% suggesting a proliferative effect (Figure 1).

**Figure 1.**
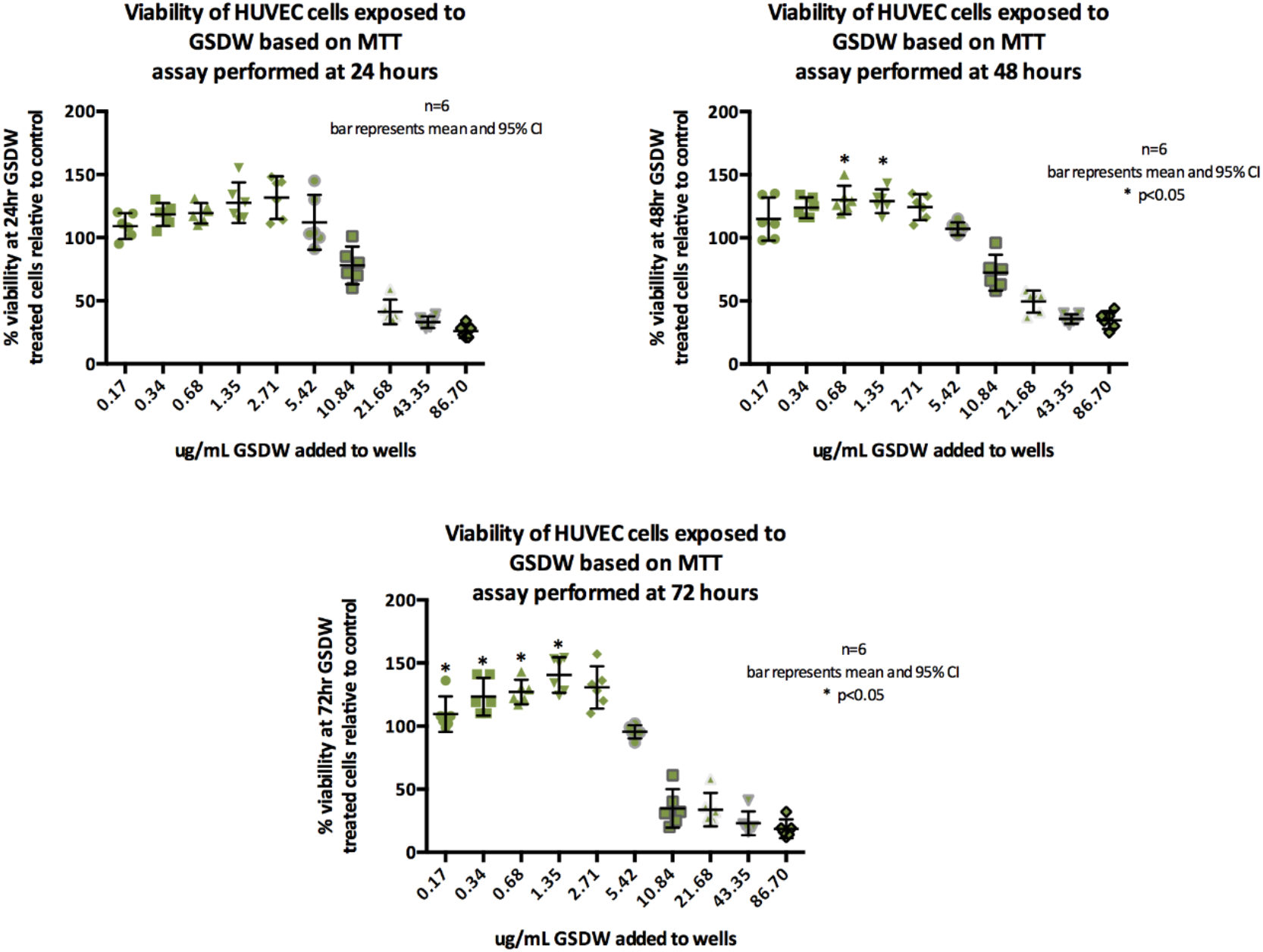
MTT Viability data for HUVECs at 24, 48 and 72.

At 24 h, the differences in cell viability from controls were not statistically significantly different according to the concentration of herb added Welch’s F (5, 13.77) = 2.066, p=0.132. At 48 h, differences between herb concentrations were statistically significant F (5, 30) = 4.398, p = 0.004, effect size is η^2^= 0.4. At 72 h, differences between herb concentrations were statistically significant F (5, 30) = 10.039, p <.0005, effect size is η^2^= 0.63. Tukey’s post hoc analysis showed that, at 48 h, viability increased above 100% when GSDW was added at concentrations of 0.68 μg/mL (95%CI 4.67 to 40.10) and 1.35 μg/mL (95%CI -.99 to 35.33). At 72 h, the difference in viability was significant with concentrations between 0.34μg/mL - 2.71 μg/mL showing greater than 100% viability; 0.34 μg/mL (95%CI −49.70 to −5.96.), 0.68 μg/mL (95%CI −53.37 to −9.63), 1.34 μg/mL (95%CI −66.97 to −23.13), 2.71μg/mL (95%CI −57.04 to −13.30).

The group means showed a statistically significantly difference (*p* < 0.05) at 48 and 72 h, indicating that viability of HUVECs at 48 and 72 h is increased by the presence of GSDW added to the cell media at concentrations of between 0.34 to 2.71 μg/mL which indicates a proliferative effect. The MultiTox assay was only performed on a single plate to confirm an increase in live cell numbers combined with a decrease in protease activity resulting from cell death suggesting that the GSDW extract is having a proliferative effect on the HUVEC cells (data not shown).

### Angiogenesis – Differentiation

In response to treatment with GSDW, HUVECs showed an increase in all four parameters of differentiation when compared to a media only control (Figure 2). No significant differences were seen for the individual herbs.

**Figure 2.**
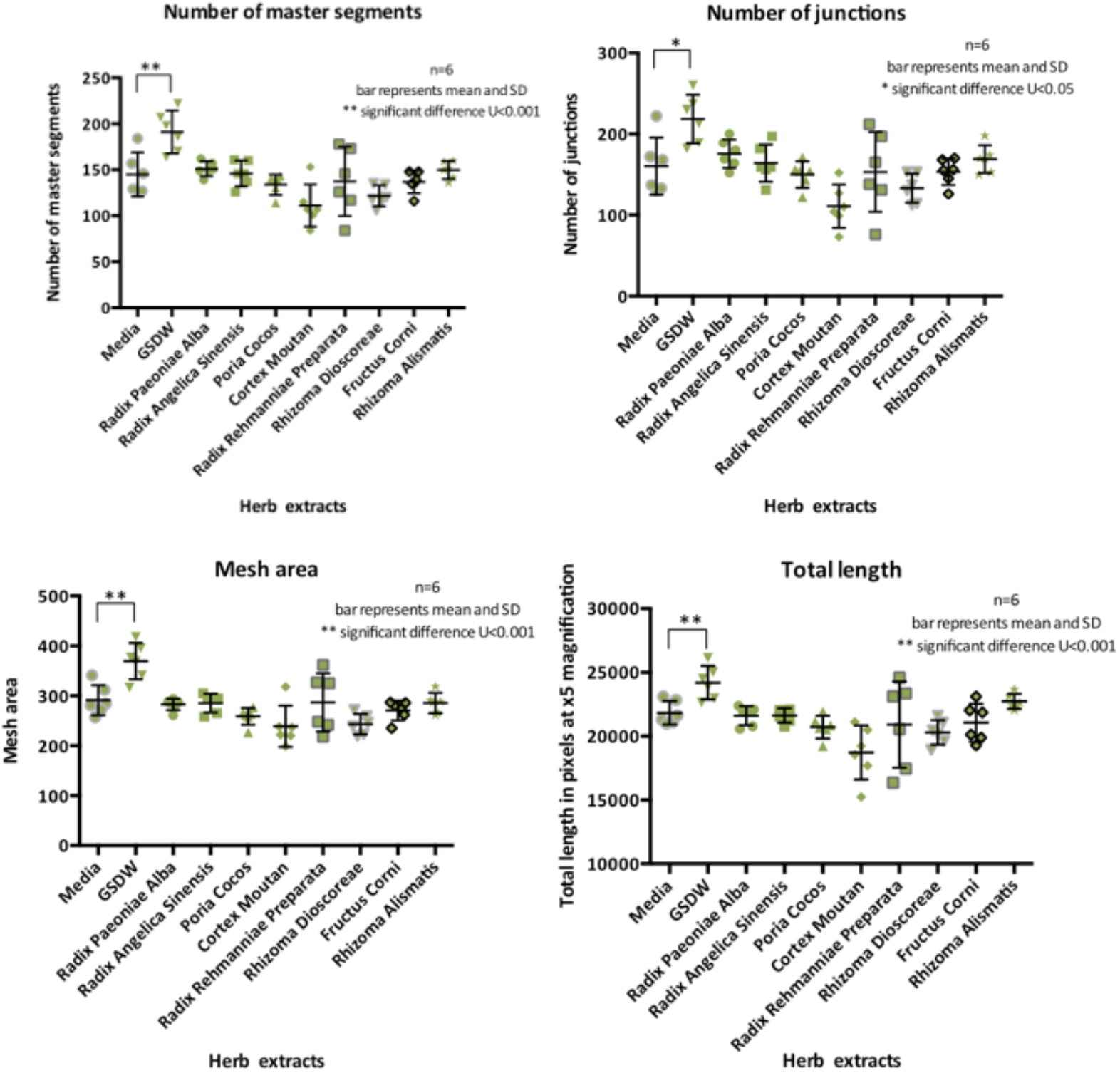
HUVEC differentiation parameters measured displayed as scatter dot plot with mean and SD.

T-tests showed statistically significant differences between GSDW and media with 0.1% FBS for number of segments 46.34±13.00 (t (10) =3.57, p=0.005), d=2.05, number of junctions 58.17±18.83 (t (10) =3.09, p=0.011) d=1.78, mesh area 78.17± 19.32(t (10) =4.05, p=0.002), d=2.33 and total length 653±910.79 (t (10) =3.62, p=0.005), d=2.09.

These observations were also seen when HUtMECs were used (Figure 3) however there were also significant differences in mesh area for all herb extracts and total length for all herb extracts except Radix Angelicae Sinensis, Sclerotium Poria Cocos and Rhizoma Dioscoreae. There was an increase in number of junctions and segments for Fructus Corni. Segment number and length was increased for Radix Paeoniae Alba, Sclerotium Poria Cocos and Rhizoma Dioscoreae and Rhizoma Alismatis.

**Figure 3.**
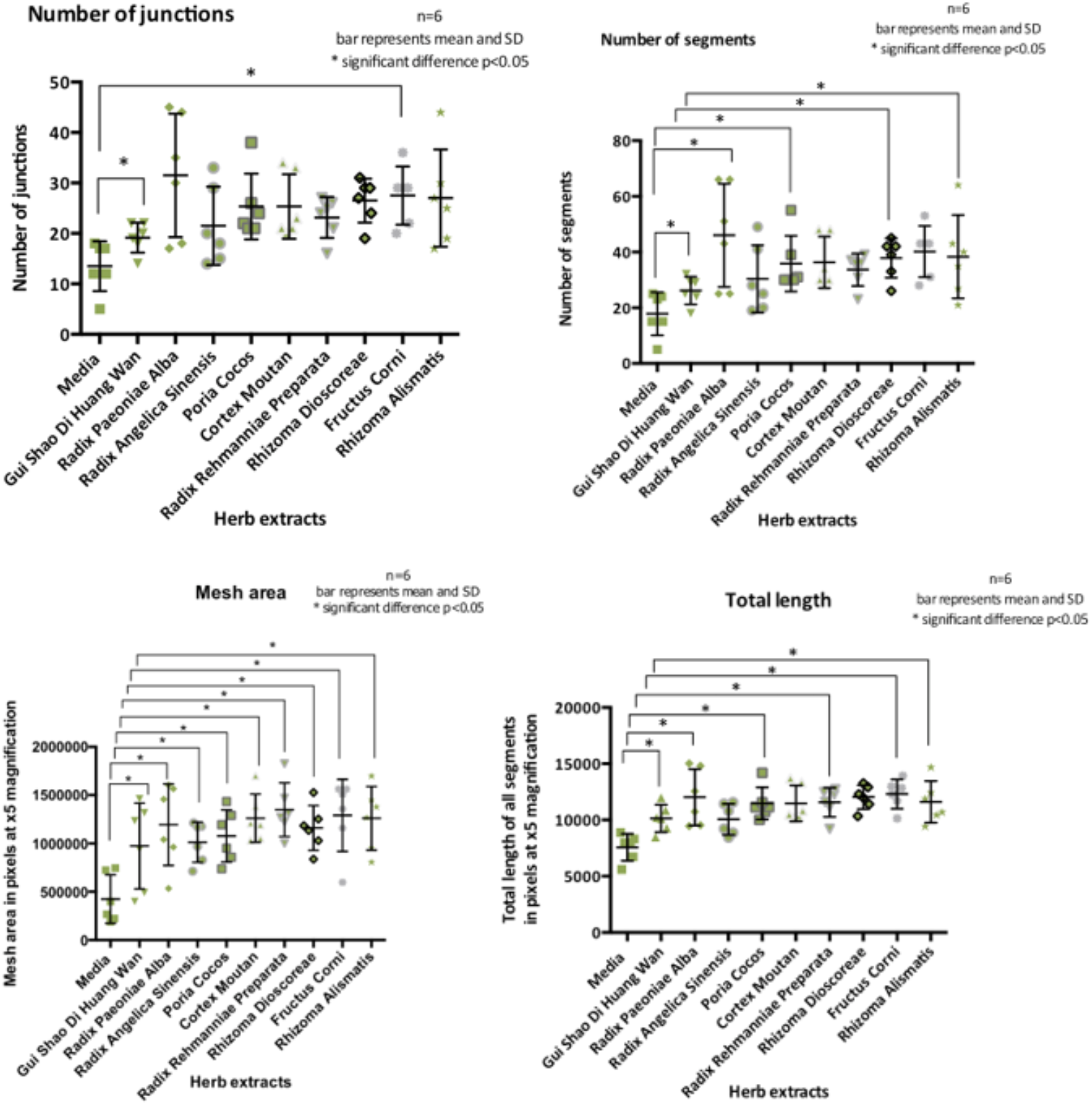
HUtMEC differentiation parameters measured displayed as scatter dot plot with mean and SD.

T-Tests showed a statistically significant difference between media and GSDW for all parameters. Data presented here as mean±SD and the effects size is Cohen’s D; number of segments 4± 1.63 (t (10) =2.46, p=0.034), d=1.42, number of junctions 5.67±2.36 (t (10) = 2.41, p=0.037 d=1.39, mesh area 549304± 208247 (t (10) =3.67, p=0.004), d=1.52 and total length 2562±697 (t (10) =2.67, p=0.004), d=2.12.

### 3.3 Evaluation of changes to oestrogen and progesterone receptor expression following GSDW treatment

Data was analysed as mean absolute volume (Fig 4 and 5). It was not possible to perform statistical analysis as there was high variability between blots. However, as confirmation, blots were also analysed as relative abundance compared to media only and normalised to a b actin loading control (data not shown) and only when there was concurrence between all three methods of analysis was a result of up or down regulation concluded.

**Figure 4.**
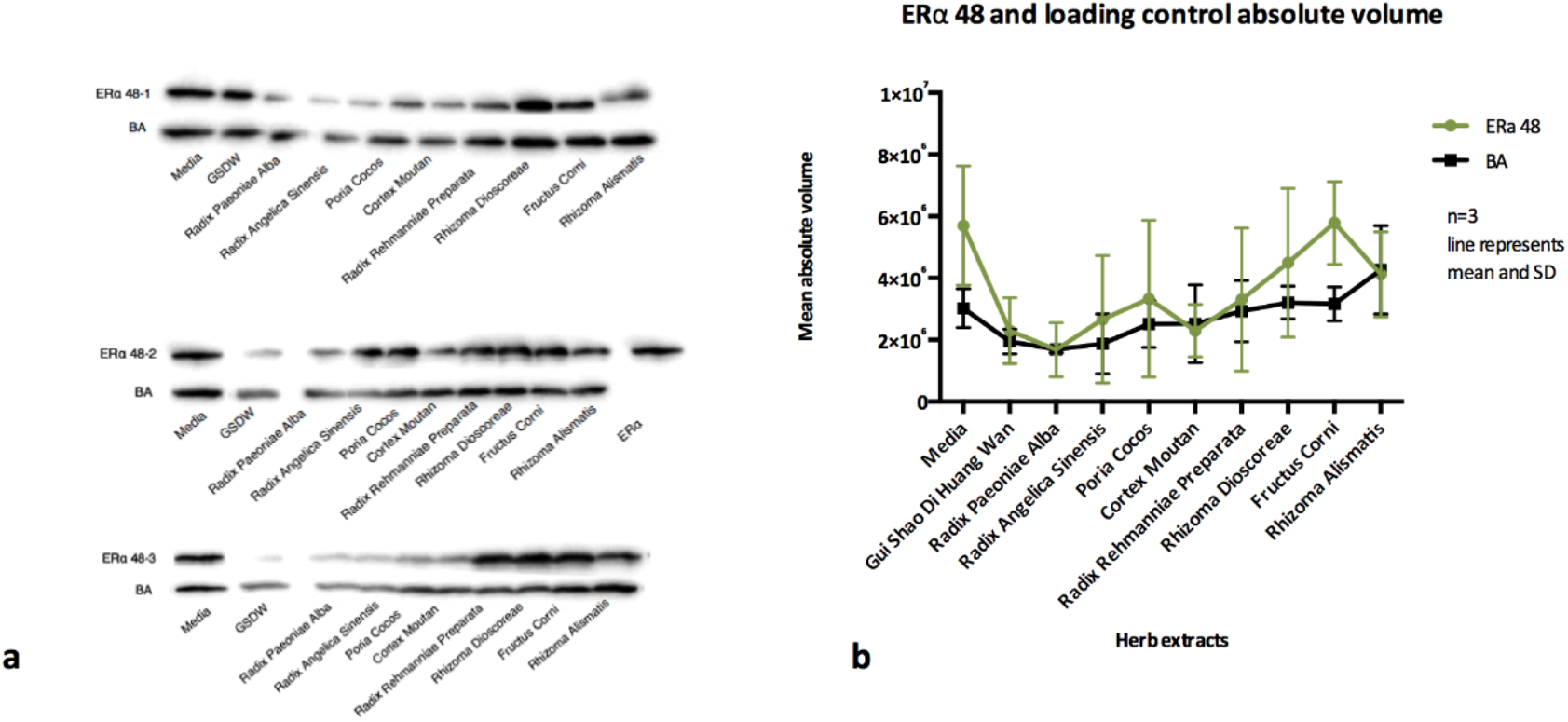
a) ERα expression from lysates of three different wells of Ishikawa cells that have been exposed to herb extracts for 48 h b) absolute volumes for ERα and βA, downregulation of ERα is shown for Radix Paeoniae Alba, Radix Angelicae Sinensis, Cortex Moutan, and Radix Rehmanniae.

**Figure 5.**
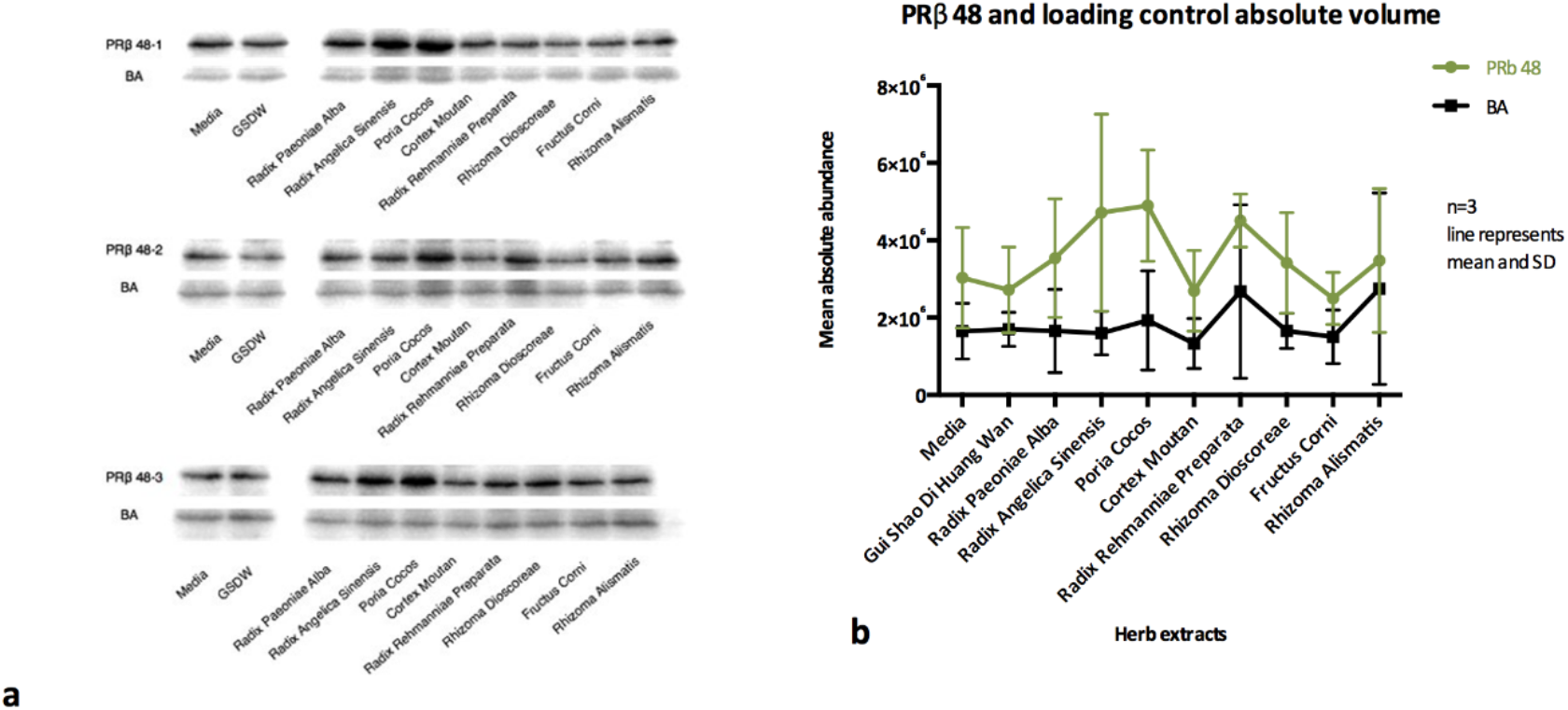
a) PRβ expression from lysates of three different wells of Ishikawa cells that have been exposed to herb extracts for 48 h b) PRβ at 48h displayed absolute volumes for PRβ and βA which shows upregulation of PRβ for Paeoniae Alba, Radix Angelicae Sinensis, Sclerotium Poria Cocos, and Radix Rehmanniae

The absolute volumes (*Figure 4b*) show downregulation of ERα in Ishikawa cells after 48 incubation with Radix Paeoniae Alba, Radix Angelicae Sinensis, Cortex Moutan and Radix Rehmanniae. This pattern of downregulation was also observed when blots were analysed by relative abundance to media only control and normalisation ratio to BA loading control (data not shown).

The absolute volumes (Figure 5b) show upregulation of PRβ in Ishikawa cells after 48 incubation with Radix Paeoniae Alba, Radix Angelicae Sinensis, Sclerotium Poria Cocos, and Radix Rehmanniae This was also seen in both the relative abundance to media only control and normalisation ratio to βA loading control (data not shown).

## 4. Discussion

The results of this study suggest that GSDW extracts, in vitro, can influence biological processes that may contribute to improved endometrial receptivity. One important factor influencing endometrial receptivity is the thickness and vascularity of the endometrium. The increased number of cells and differentiation into pseudo tubules could explain these changes. Modification to the sex steroid receptors will influence the response to endogenous and exogenous sex steroids, oestrogen and progesterone.

Proliferation was investigated using an MTT assay and HUVECs. After menstruation, the epithelial cells of the endometrium proliferate in order to repopulate the shed functionalis. If epithelial cells were to proliferate without concurrent proliferation of endothelial cells a thickening of the endometrium with poor uterine perfusion could occur, which would have poor prognosis for implantation. This study showed increased viability indicative of proliferation, verified in the MultiTox assay. The herb extract was able to promote cell proliferation at 48 h from 0.68 to 1.35 μg/mL and at 72 h from 0.34 to 2.71 μg/mL.

Although HUVECs are a useful model they are not the same as the cells that are involved in implantation. Endothelial cells in the umbilical vein (HUVEC) may be continuous and they are polygonal because fluid exchange occurs at the placenta not in the veins themselves. The uterine HUtMECs are likely to be fenestrated as the myometrium where the cells originate is a tissue where fluid exchange is occurring ^28^. The HUtMECs are microvascular cells from the myometrium, which are one of the multiple tissue types with which an implanting blastocyst will interact. It will first come into contact the luminal epithelium, displacing this to invade the basal lamina and into the stroma ^29^. The myometrium where HUtMECs originate is below the stroma. It is not the tissue most dynamically responsive to the sex steroids, nor is it part of the regenerative basal layer. It is however, a closer tissue type to those intimately involved in implantation than HUVECs, and the vascularity of the myometrium will affect the downstream uterine perfusion and therefore delivery of nutrients, these may be sufficiently similar to be a useful model.

This study showed that metrics of differentiation were increased in both HUVEC and HUtMEC cells in response to the herb extracts. These cells showed significant differences in differentiation when exposed to GSDW than a media only control for all four measured parameters of differentiation, number of segments, number of junctions, length of tubules and mesh area. These effects on the tubules suggests that there are compounds within the herb extracts promoting differentiation. It is unclear exactly how these metrics translate into changes in the endometrium. Is an increase in length advantageous? Does it lead to greater penetration of the myometrium? Translating the metrics of tubule development to function of the vessels *in vivo* is not direct. The formation of a network, it is argued is not the same process as a functional lumen and therefore a blood containing vessel ^30^. A functional lumen is in part stimulated to be formed by blood flow ^31^. A matrigel assay is only a 2D representation of a complex 3D process.

Ishikawa cells are immortalized epithelial cells and are clones of a human endometrial adenocarcinoma and they have been shown to express both oestrogen and progesterone receptors. There was a downregulation in ERα by Radix Paeoniae Alba, Radix Angelicae Sinensis, Cortex Moutan and Radix Rehmanniae and upregulation of PRβ for Radix Paeoniae Alba, Radix Angelicae Sinensis, Sclerotium Poria Cocos, and Radix Rehmanniae at 48 h. It would appear that the steroid receptors ERα and PRβ are affected by the treatment but whether this is advantageous to receptivity or not would depend on the stage of the cycle. Downregulation of ERα would be beneficially early in the cycle and again from the time of ovulation and through to implantation, but needs to be upregulated for migration and proliferation during the secretory phase ^32^. Upregulation of PRβ would support the changes required for differentiation of the vessels ^33^, implantation ^6^ and maintenance ^34^ of the pregnancy, but would be inhibitory to the migration and differentiation during the middle part of the secretory phase. In specific disease processes these actions may be of benefit. Downregulation of ERα may be appropriate when treating luteal phase defects and polycystic ovarian syndrome ^35^. Upregulation of PRβ could be appropriate in endometriosis ^19^. All *in vitro* cellular assays, even with the primary cell lines are in an artificial environment, without the normal stimulation of their environment and neighbouring cells, different epigenetic changes will occur. Endothelial cells forming structures will behave differently in contact with pericytes and smooth muscles cells and subject to the sheer stress of vascular flow ^36^.

A non cancer cell line would be more representative and normal endometrial cells can be cultured from patient derived samples or bought as endometrial tissue blocks commercially. Research can be performed on biopsied cells but these are phenotypically unstable and do not grow well. Few cell lines were available to purchase other than cancer cell lines, such as the Ishikawa’s which are epithelial-like but there are transformed human endometrial, epithelial, and stromal cells. Repeating the assays with a wider range of cells types might yield further interesting results that would show effects in a wider representation of the uterine tissues involved in receptivity.

*In vitro* cell assays have significant limitations and cannot be taken as direct evidence that the same behaviour would exist in the specific cell type *in vivo.* When looking at therapeutic effects it may be necessary to be able to model the actual pathology of the disease to see true effects. These assays do allow investigation of specific functions outside the uncontrolled variables of the organism and so have their place in a diverse portfolio of evidence.

Another limitation of using a cellular assay is that the cells in the experiments are presented with the whole extract, not a selection of compounds which may have been altered when absorbed through the gut as would occur in a patient. When a person consumes a herbal medicine, there will be food in their digestive tract, they will each have a unique microbiome that modifies absorption, their own metabolome, liver enzymes, and body volume. Pharmacokinetic studies show that ingestion of a substance is not the same as absorption and a compound may be modified during absorption. There can be up to 60 types of bacteria in the colon which catalyse drug reactions. Initial metabolism can occur in the intestines such as oxidation, reduction, hydrolysis as well as glucorination, sulphation, *N*-acetylation, methylation, gluthianone and glycine conjugation. Once through the gut wall it must then pass through the liver where substances are subject to first pass hepatic metabolism which modifies many compounds. Food changes the absorption of drugs and distribution will depend on absorption, permeability, diffusion, perfusion and tissue binding ^37^.

## 5. Conclusions

The *in vitro* assays performed indicate that aqueous extracts of GSDW have the potential to influence factors determining the vascularity of the endometrium in that they increased both the proliferation of endothelial cells and parameters of endothelial cell differentiation, as summarised in Figure 6. Furthermore, these functions are also in part regulated by the ovarian steroid hormones and the expression of these receptors are also seen to be modified by extracts of GSDW.

**Figure 6.**
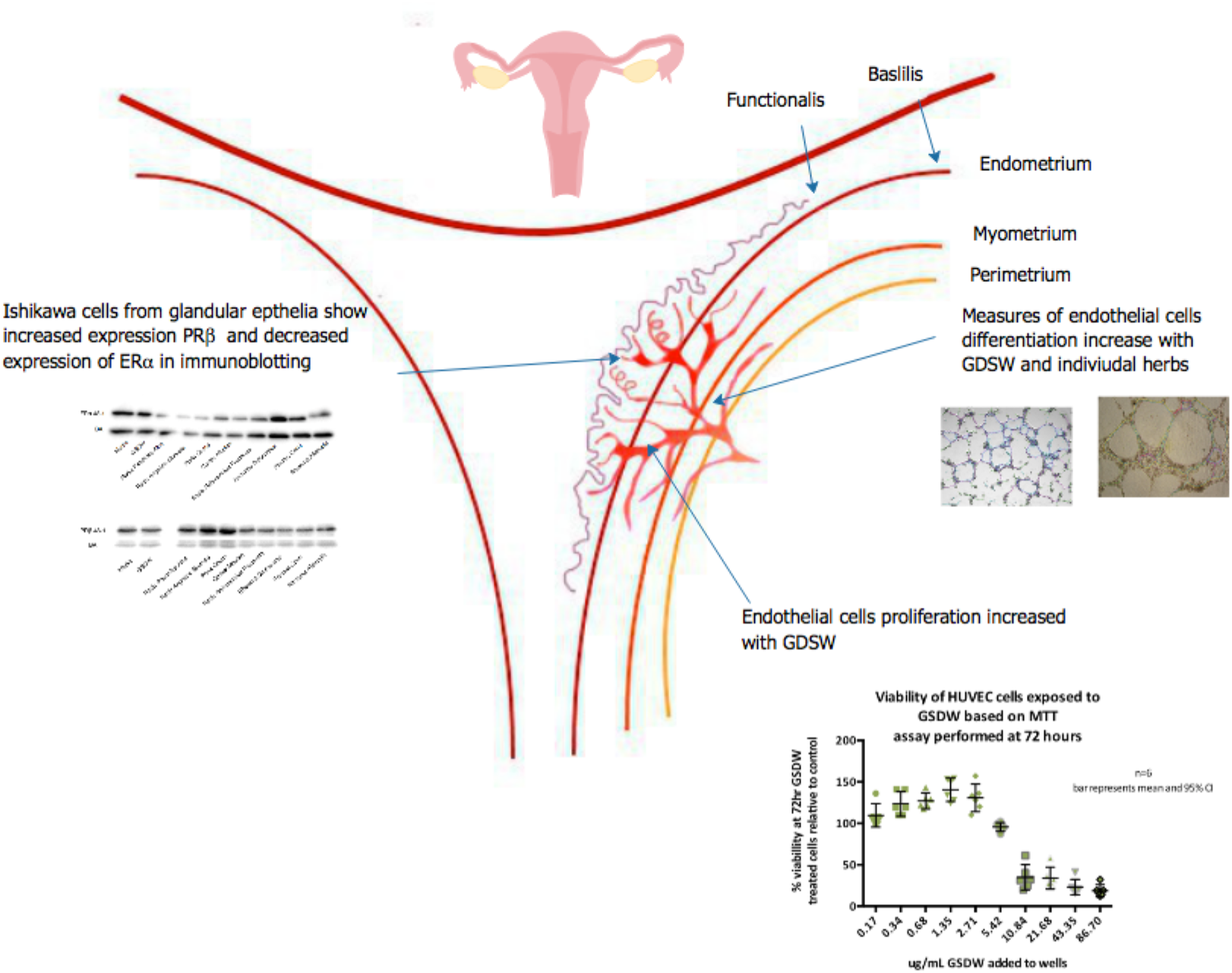
Summary of research presented on the effects of GSDW on angiogenesis proliferation, differentiation, and Immunoblotting assays.

